# Transcranial Pulse Stimulation with Ultrasound in Alzheimer’s disease – A new navigated focal brain therapy

**DOI:** 10.1101/665471

**Authors:** R. Beisteiner, E. Matt, C. Fan, H. Baldysiak, M. Schönfeld, T. Philippi Novak, A. Amini, T. Aslan, R. Reinecke, J. Lehrner, A. Weber, U. Reime, C. Goldenstedt, E. Marlinghaus, M. Hallett, H. Lohse-Busch

## Abstract

Ultrasound-based brain stimulation techniques offer an exciting potential to modulate the human brain in a highly focal and precisely targeted manner. However, for clinical applications the current techniques have to be further developed. We introduce a new ultrasound stimulation technique, based on single ultrashort ultrasound pulses (transcranial pulse stimulation, TPS) and describe a first navigable clinical TPS system. Feasibility, safety and preliminary (uncontrolled) efficacy data in Alzheimer’s disease (AD) are provided. Simulation data, *in vitro* measurements with rat and human skulls/brains and clinical data in 35 AD patients were acquired in a multicentric setting (including CERAD scores and functional MRI). Preclinical results show large safety margins and patient results show high treatment tolerability. Neuropsychological scores improved significantly when tested immediately as well as 1 and 3 months after stimulation and fMRI data displayed significant connectivity increases within the memory network. The results encourage broad neuroscientific application and translation of the new method to clinical therapy and randomized sham-controlled studies.

## 1. Introduction

Recently, several publications have reported the potential of ultrasound to stimulate the human brain in a highly focal and precisely targeted manner. However, for clinical applications the current techniques have to be further developed and certified mobile systems for clinical research and application are not yet available. Here we introduce a new ultrasound stimulation technique, which was specifically developed for clinical applications by an interdisciplinary consortium for brain stimulation (MH), clinical neuroscience (RB), clinical ultrasound (HLB) and ultrasound technology (EMar). The new method is based on single ultrashort ultrasound pulses (transcranial pulse stimulation, TPS) that avoid secondary stimulation maxima or brain heating (Mueller et al. 2017). In contrast to electrophysiological brain stimulation techniques that often suffer from conductivity effects (Minjoli et al. 2017) and lack of deep stimulation capabilities (Spagnolo 2018), the target for ultrasound-based neuromodulation can be spatially distinct, highly focal, and is not restricted to superficial layers of the brain. For brain therapy, this enables a controlled modulation of a specific brain region with reduced probability of producing unwanted co-stimulations of other brain areas. Clearly defining which brain areas are affected by the stimulation and which are not, is an important advance for clinical application and neuroscientific research. Previous studies have already demonstrated that non-invasive focal ultrasound applications may modulate the function of healthy human brains (e.g., Legon et al. 2014, 2018, Lee et al. 2015). Typically, these are ultra short term neuromodulations but also neuromodulation effects of more than one hour have recently been shown in macaque monkeys (Verhagen et al. 2019). Initial clinical data concerning non-navigated stimulation (Lohse-Busch et al. 2014) and a case report (Monti et al. 2016) are also available.

Currently, two principles for ultrasound based focal neuromodulation of human brains with mobile single ultrasound transducers exist: (1) Application of ultrasound beams consisting of 300-500 ms trains of ultrasound waves (<0.7 MHz) repeated every 3-6 s (Low-intensity Focused Ultrasound technique, LIFUS). (2) Application of single ultrashort (3 μs) ultrasound pulses with repetitions every 200-300 ms (TPS, Figure 1). The principal technology used for TPS has the advantage that it represents a modification of a technology already in long-term clinical use for standard clinical treatments. Standard indications comprise neurology (e.g., spastic muscle paralysis), cardiology (e.g. ischemic heart disease), orthopedics (e.g. fasciitis, tendinopathy), dermatology (e.g., wound healing) and urology (e.g., vascular erectile dysfunction). TPS represents a new extension of this technology and is characterized by the application of short pulses which are dominated by lower frequencies to improve skull penetration.

**Fig. 1.**
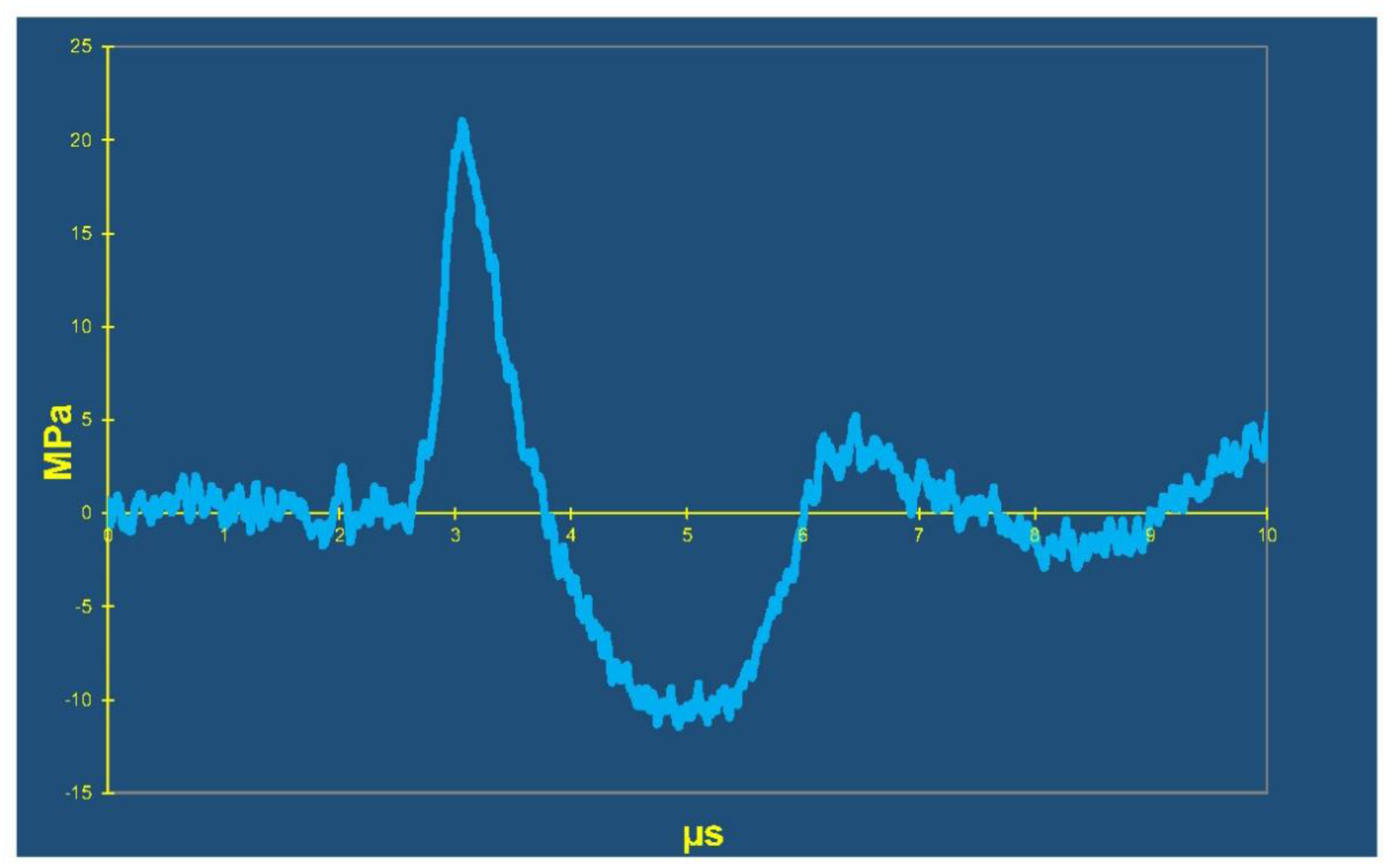
TPS pulse wave characteristics. TPS pulse wave with about 3 μs duration (MPa = Mega Pascal). In contrast, the typical duration of a “pulse” generated by Low Intensity Focused Ultrasound (LIFUS) techniques is 300-500ms, consisting of an ultrasound train in the low MHz range (<0.7 MHz). For the clinical study we applied 6000 TPS pulses per therapeutic session. Repetition frequency was 5 Hz and pulses were distributed over various brain regions.

In this manuscript, we describe a first navigable clinical TPS system and provide feasibility and safety data in the context of Alzheimer’s disease (AD, Figure 2). The feasibility for non-invasive targeted and focal energy deposition was tested by simulations and in vitro measurements with rat and human skulls as well as brain specimens. This was followed by a multicenter study on feasibility, clinical safety and preliminary efficacy. Since we expect TPS to be applied as an independent add-on therapy, we only included patients already receiving optimized standard clinical treatment. Preliminary efficacy was evaluated based on the neuropsychological CERAD (Consortium to Establish a Registry for Alzheimer’s disease, Ehrensperger et al. 2010) test battery and associated scores as primary outcomes. Depression monitoring was done with the Geriatric depressions scale (GDS) and the Beck Depression Inventory (BDI). As secondary outcome, high field fMRI data were recorded in a subgroup of 19 patients. For safety evaluation, clinical patient investigations, patient questionnaires and MR images were used.

**Fig. 2.**
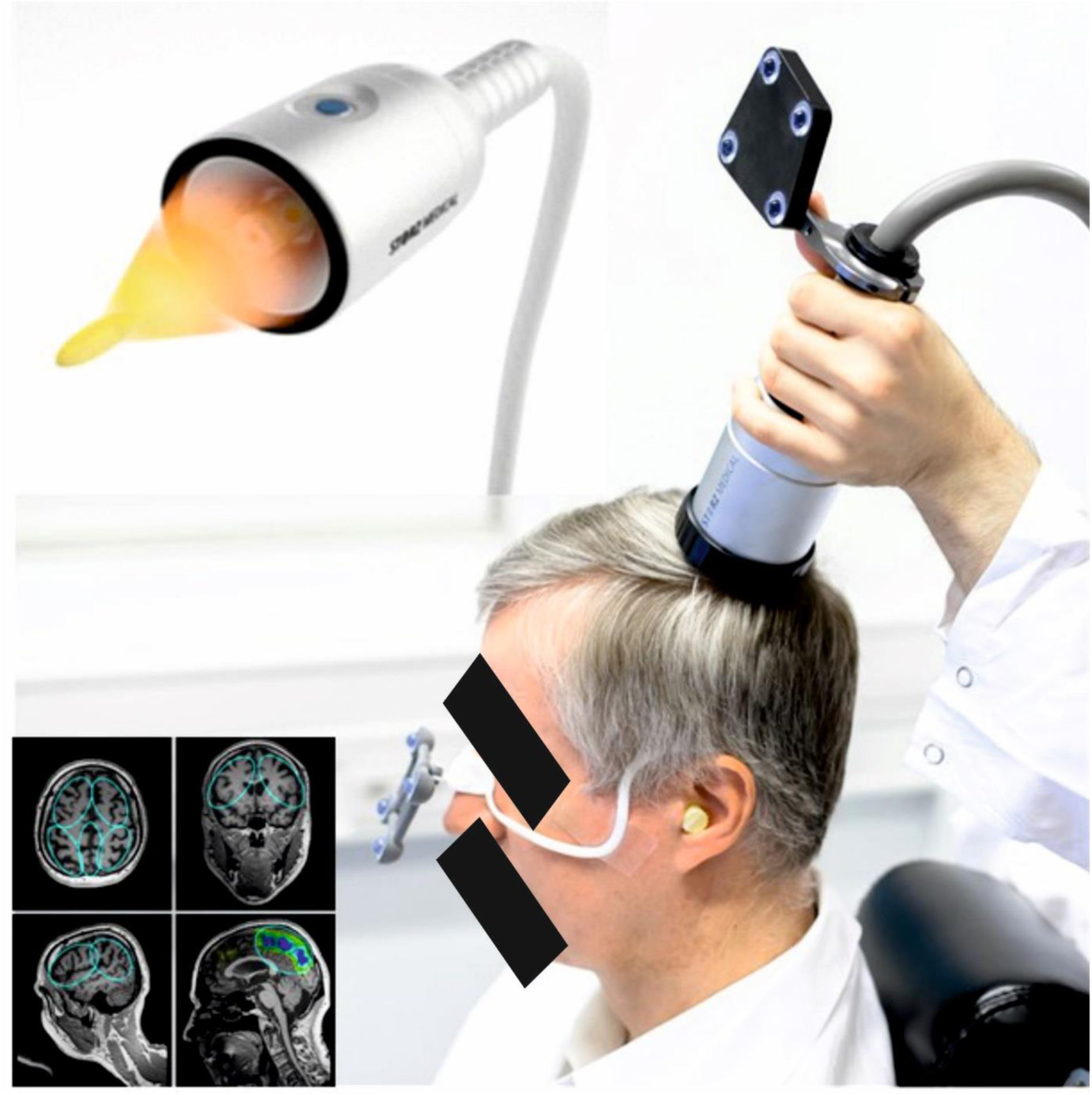
Clinical setting for navigated TPS therapy. Goggles and TPS handpiece are equipped with infrared reflectors for visualization and tracking of the TPS focus (which is illustrated in the left upper insert) to regions of interest in individual anatomical MR images (blue circles, left lower insert).

## 2. Methods

### 2.1. TPS system

The newly developed TPS system consists of a mobile single transducer and an infrared camera system for MR based neuronavigation (NEUROLITH, Storz Medical AG, Tägerwilen, Switzerland, Figure 2). TPS generates single ultrashort (3 μs) ultrasound pulses with typical energy levels of 0.2-0.3 mJ/mm2 and pulse frequencies of 1-5 Hz (pulses per second). Stimulation of target areas is done via variable stand offs at the handpiece for depth regulation and manual movement of the handpiece over the skull with immediate visualization of the individual pulses on the patients’ MR images. For highly focal applications, the handpiece may be fixed at a constant position over the skull. The whole treatment session can be recorded for post hoc evaluation of the individual intracerebral pulse localizations. All procedures are compliant with all relevant ethical regulations and all study procedures were approved by the responsible authorities. For the clinical study all patients gave their written informed consent.

### 2.2. TPS focal energy transmission

The simulations and in vitro experiments were set up to investigate the following issues: (1) Can TPS transmit energy adequately through the skull? (2) Can TPS generate a small focused stimulation beam below the skull?

#### 2.2.1. TPS data simulations

Temporal peak intensity fields as generated by the clinically applied TPS system have been simulated for free degassed water and two real skulls including brain tissue. The numerical models were reconstructed from the CT scans of the complete heads of two donors. The position of the TPS source in relation to the skull was recreated, according to the configuration of the experimental measurements described below. The numerical simulations were performed using Matlab (Mathworks, USA) and the open-source k-Wave toolbox, which uses a k-space pseudo-spectral time domain solution to coupled first-order acoustic equations (Treeby and Cox 2010). The simulation was limited to a volume (98×50×50 mm³) of the head containing the expected focal area and the surroundings (Figure 3) as extracted from the CT scans. The Hounsfield Units were converted into density and acoustic celerity using the built-in k-wave functions based on the empirical results of Schneider et al. (1996) and Mast (2000). Absorption coefficients of 3.57 dB.cm^-1^.MHz^-1^ and 0.58 dB.cm^-1^.MHz^-1^ were respectively assigned to bone and brain structures (Szabo 2014). The region outside of the skulls was modeled as non-absorbing water (ρ = 1000 kg.m^-3^, c = 1489 m.s^-1^). The non-linearity parameter B/A was set to 7, corresponding to most of biological soft tissues including brain (Szabo 2014, Hamilton and Blackstock 1998), for the whole computational domain. The pressure source was modeled as a brass parabolic reflector (c = 4198 m.s^-1^; ρ = 8470 kg.m^-3^) centered on a cylindrical coil, matching dimensions those of the real device. The initial acoustic excitation was simulated as a cylindrical pressure wave uniformly distributed over the coil and modelled as a single-pulse.

**Fig. 3.**
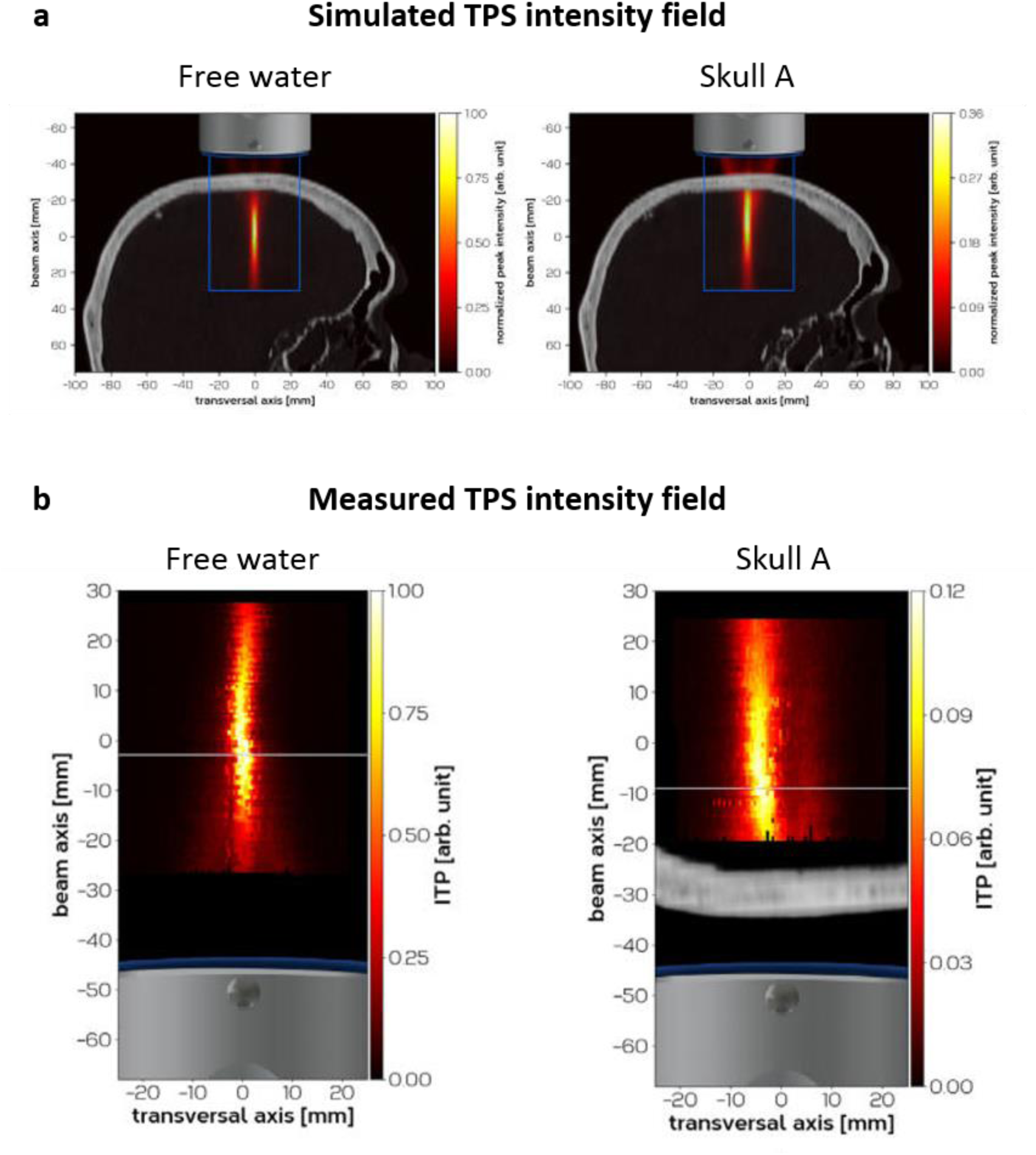
Simulated and measured TPS intensity field. **a,** Simulated temporal-peak intensity field for the simulation for free water (left, but skull overlaid) and for skull A (right). The blue lines indicate the limits of the computational domain. Note skull attenuation of about 65% as indicated by the color bar. **b**, Measured temporal-peak intensities for free water (left) and for skull A (right). The white line indicates the spatiotemporal peak (beam axis maximum). Note that skull attenuation is even stronger (about 85%) than expected from the simulations.

#### 2.2.2. TPS in vitro measurements

##### Human skull and brain samples

Single pulse pressure waves were generated by a device with the same acoustic performance as the system used for the clinical study (Figure 2, Storz Medical AG, Tägerwilen, Switzerland).. A typical pressure pulse generated by this device, measured at the focus, is shown in Figure 1 and the experimental setup is illustrated in Figure 4. The pressure pulses were measured using a needle hydrophone (Dr. Müller Instruments, Oberursel, Germany) fixed on a two-axis sliding stage. The predefined measurement domain (50 mm along the beam axis and 40 mm along the transversal axis) was centered on the geometrical focus of the handpiece, defined as the origin of the coordinate system. The spatial transversal and axial measurement steps were kept below 1 mm and 3 mm respectively. All pressure waves were released at a drive level of 0.25 mJ/mm^2^, and at a pulse repetition frequency of 2 Hz. First, a reference acoustic field measurement was performed in free water. Then, a section of human skull (roughly intermediate between bregma and lambda), with the brain parenchyma completely removed, was placed in front of the handpiece and firmly fastened in a holder. The relative position of the handpiece to the skulls was determined from photographic acquisitions during the measurements. A 3D reconstruction of the TPS handpiece and the mounting plate of the water basin was recreated in Blender software [https://www.blender.org/]. The CT scans were segmented to create a 3D surface model of the human samples. Virtual cameras using the specifications of the Canon EOS 5D Mk II were then aligned and positioned to match the reference images. The plane of incision around the circumference of the skull was determined to create a 3D model of the skull section, which was then used to reconstruct the measurement setup. These steps allowed a discrete transformation between the CT image system and real world geometric focus position. The CT data was transformed and interpolated to the resolution of 200um, which allowed for an easy extraction of both the 3D computational volume for the simulations described above and the image plane for the visualization of the 2D measurements. The measured fields were displayed in the corresponding slice in such a way that the origin of the coordinate system of the measurement setup, representing the geometrical focus, matches the corresponding position in the CT slice (Figure 3). A similar procedure was used for measuring the pressure drop in 10 human brain samples in vitro (soft tissue without skull, 0-7 days post mortem).

**Fig. 4.**
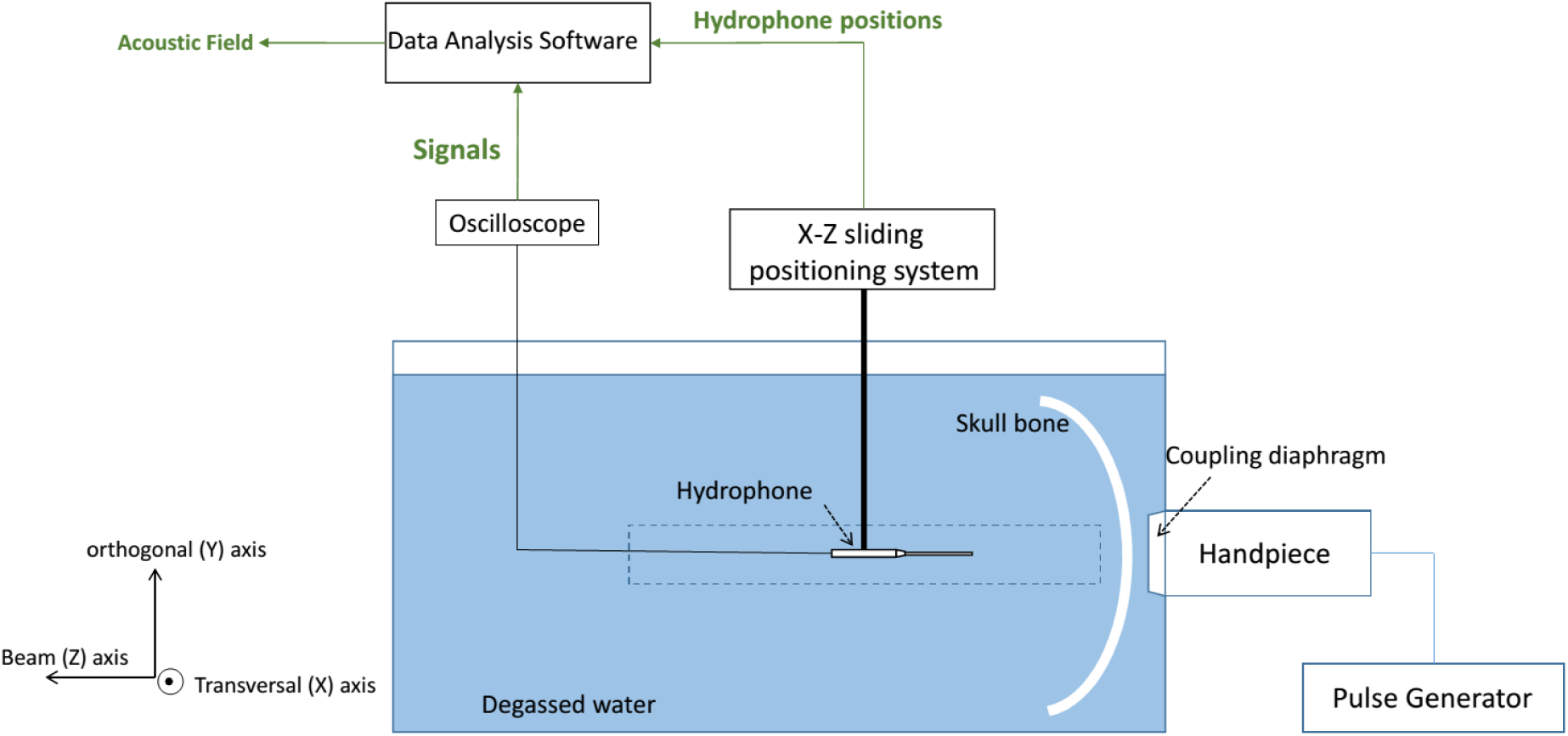
Experimental setup for the in vitro measurements. The TPS handpiece was fixed on the side of a basin filled with degassed water. Test specimens (e.g. human skulls, rat skulls, brain specimens) were fixed immediately in front of the handpiece. Pulse attenuations due to the specimens were recorded by the Hydrophone with reference to free water results (see Figure 3 and 6). The Hydrophone can be moved in 3D.

##### Rat Skull

Allowing judgements about expectable differences between animal skulls and human skulls, we also performed measurements of a rat skull with the same principal technology. For this, TPS was applied with 0.1 mJ/mm^2^, 0.35 mJ/mm^2^ and 0.55 mJ/mm^2^ and at 6 positions of the rat skull: bregma point, about 5 mm left and right from the bregma, lambda point and about 5 mm left and right from the lambda (Figure 5). Pulse amplitude was measured at the TPS focus below the skull.

**Fig. 5.**
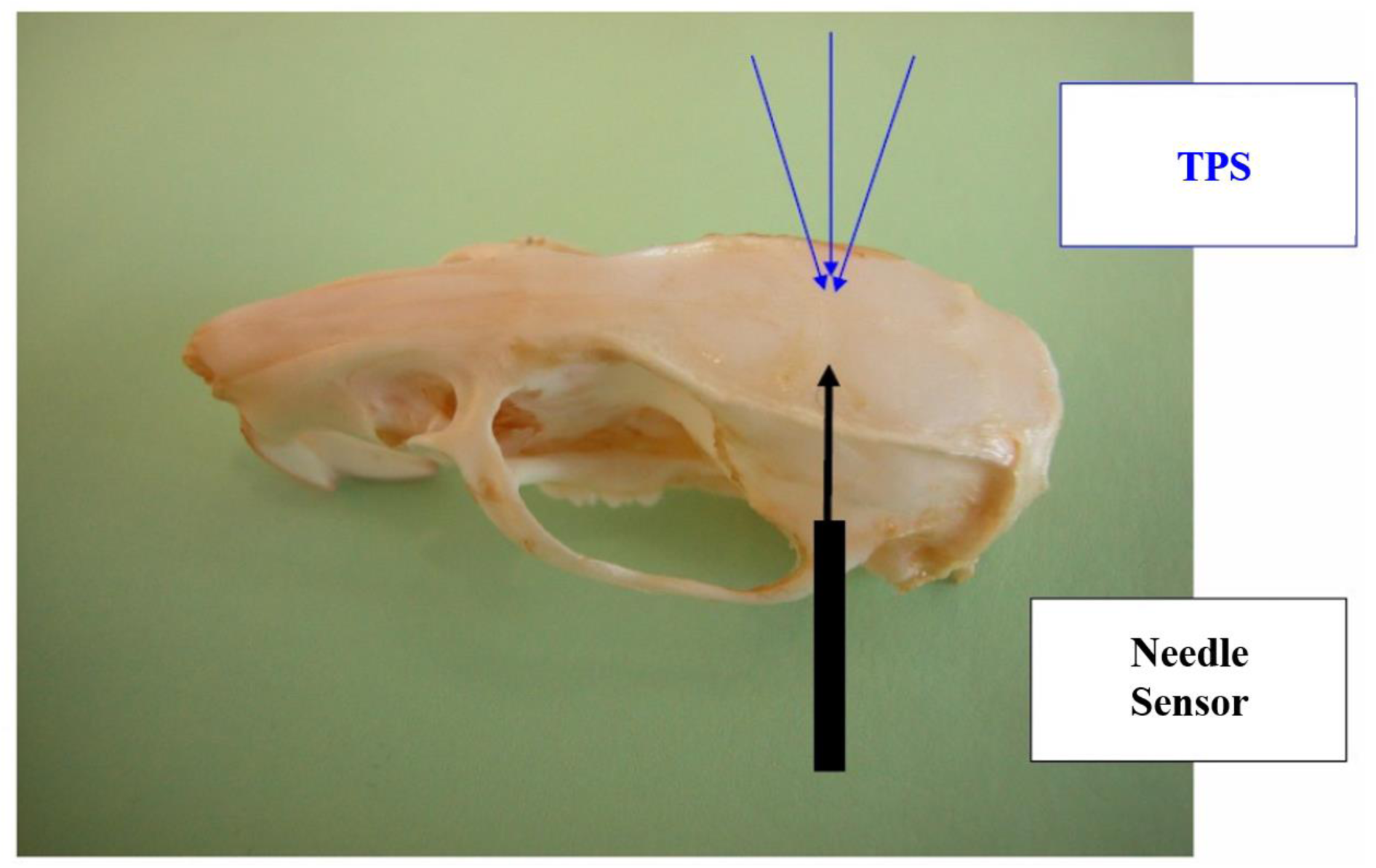
Setup of rat skull measurements. TPS pulses with the TPS stimulator were applied through various locations of the rat skull. Attenuation of the pulse intensity was measured at peak pulse intensity below the skull with a needle sensor (Müller-Platte needle hydrophone, Dr. Müller Instruments, Oberursel, Germany). On average, we measured only 1/3 of the attenuation measured with human skulls. This illustrates the importance of comparative measurements when trying to translate new ultrasound techniques to human applications.

### 2.3. TPS safety investigations in anesthetized rats

Single pulse pressure waves were generated by a device equivalent to the system in the clinical study (Storz Medical AG, Tägerwilen, Switzerland). 80 male Sprague-Dawley rats (Charles-River, Germany) were treated with TPS applied at a fixed position over the rat skull and a constant focus in the brain under anesthesia (Ameln-Mayrhofer et al., in preparation). Stimulation was performed at a frequency of 3 Hz in groups of 10 rats each −1 control and 7 test groups with the following settings: 0.1 mJ/mm2, 100 pulses; 0.1 mJ/mm2, 200 pulses; 0.1 mJ/mm2, 400 pulses; 0.2 mJ/mm2, 100 pulses; 0.2 mJ/mm2, 200 pulses; 0.2 mJ/mm2, 400 pulses; 0.3 mJ/mm2, 100 pulses. For safety evaluations, brain preparations (80 rats) and histological investigations (16 rats) were performed to investigate for possible intracerebral bleeding and tissue damage as primary outcomes. Outside the safety context of this study, animal behavior was also analysed (Ameln-Mayrhofer et al., in preparation). In more detail, rats were held in groups in Makrolon-IV cages at fixed climatization and 12h:12h light-darkness cycles. For anesthesia isoflurane 1-2% and fentanyl 5μg/kg or Butorphanol 3,3 mg/kg were used. Analgesia was required for controlled experimental conditions and a stable brain stimulation focus. For postsurgical analgesia carprofen was used. For post treatment brain preparations, animals were decapitated.

### 2.4. TPS multicenter Alzheimer’s disease study

The multicenter clinical pilot study was designed to investigate the following issues: (1) Is TPS safe and feasible in a broad range of patients and with varying treatment durations? (2) Are there indications for preliminary effects as investigated by neuropsychological scores and fMRI data? (3) Does the mode of TPS application show relevant differences concerning issues (1) and (2)? For the latter we compared a non-navigated global cortical stimulation (center 2) with a region of interest (ROI) based stimulation (center 1) with precise targeting of cortical AD network areas and clearly defined stimulation borders (requiring high focality of the technique).

#### 2.4.1. Patients

To adequately evaluate safety and feasibility on a wide range of patients, our TPS pilot study was performed within a broad clinical setting for outpatients with memory complaints related to probable AD. Although most patients suffered from mild to moderate AD (Mini-Mental State Examination (MMSE) value ≥ 18), we did not use a MMSE cutoff to minimize inclusion criteria and to enable patient variability (including controlled and stable comorbidities, see Tables 1, 2). Relevant intracerebral pathologies unrelated to AD and independent neuropsychiatric disease (like preexisting depression) were excluded. One center in Austria (center 1 Vienna, lead) and one center in Germany (center 2 Bad Krozingen) included 20 AD patients each. Only patients receiving optimized standard treatments were accepted and inclusion was based on clinical judgement and external clinical MRIs. Recruitment was performed by independent neurologists with consecutive referrals to the study centers. Due to dropouts, per protocol analysis was possible for 35 patients (19 Center 1, 16 Center 2).

**Table 1.** Demographic details and clinical baseline characteristics of the patient groups.

|  |  | **Both Centers**<br>N = 35<br>(20 female) | **Center 1**<br>N= 19<br>(12 female) | **Center 2**<br>N = 16<br>(8 female) | **Difference**<br>**between**<br>**Centers\*** |
| --- | --- | --- | --- | --- | --- |
| **Age** | Mean (SD) | 70.37 (8.57) | 68.68 (9.49) | 72.38 (7.11) | *P* = .209 |
|  | Range | [51-84] | [51-84] | [58-84] |  |
| **Education** | Mean (SD) | 11.60 (3.60) | 10.00 (2.00) | 13.50 (4.18) | *P* = .007 |
|  | Range | [7 - 20] | [7 - 14] | [8 - 20] |  |
| **Baseline MMSE** | Mean (SD) | 20.97 (5.86) | 19.53 (6.92) | 22.69 (3.81) | *P* = .113 |
|  | Range | [3 - 30] | [3 - 30] | [17 - 29] |  |
| **Baseline BDI** | N | 29 | 14 | 15 | *P* = .063 |
|  | Mean (SD) | 5.62 (5.34) | 7.36 (5.09) | 4.00 (5.21) |  |
|  | Range | [0 - 18] | [0 - 16] | [0 - 18] |  |
| **Baseline GDS** | N | 33 | 18 | 15 | *P* = .145 |
|  | Mean (SD) | 3.00 (2.47) | 3.50 (2.50) | 2.40 (2.38) |  |
|  | Range | [0 - 10] | [0 - 9] | [0 - 10] |  |

\*Differences between centers regarding demographic and clinical characteristics were tested using unpaired T-tests (two-sided, $\alpha = 0.05$ ) for normally distributed variables (age, baseline MMSE) and non-parametric Mann-Whitney-tests (two-sided, $\alpha = 0.05$ ) for education, baseline Beck Depression Inventory score (BDI), and baseline Geriatric Depression Score (GDS). SD = Standard deviation, MMSE = Mini-Mental State Examination

**Table 2.** Individual patient characteristics for Center 1 (C1) and Center 2 (C2)

| **Patient ID** | **Age** | **Gender** | **Education** | **Baseline MMSE** | **Comorbidities** |
| --- | --- | --- | --- | --- | --- |
| C1_01 | 64 | female | 11 | 3 | Suspected epilepsy, tendency to collapse, hyperlipidemia, osteoporosis |
| C1_02 | 68 | female | 8 | 12 | Hypertension, hypertensive encephalopathy |
| C1_03 | 72 | female | 11 | 23 | Vitamin B12 deficiency |
| C1_04 | 71 | male | 11 | 14 | Parkinsonian syndrome, depressive adjustment disorder, hypertension |
| C1_05 | 75 | male | 9 | 18 | AD associated depression |
| C1_06 | 82 | female | 7 | 20 | None |
| C1_07 | 81 | female | 7 | 17 | None |
| C1_08 | 76 | female | 12 | 18 | None |
| C1_09 | 56 | male | 9 | 26 | AD associated depression, panic disorder, adjustment disorder, gastritis, hypertension |
| C1_10 | 67 | male | 14 | 26 | Recurrent AD associated depression, vascular MAP, arachnoid cyst |
| C1_11 | 60 | female | 8 | 24 | Panic attacks |
| C1_12 | 84 | female | 8 | 24 | None |
| C1_13 | 61 | male | 12 | 25 | None |
| C1_14 | 61 | female | 10 | 6 | None |
| C1_15 | 75 | female | 8 | 21 | Vascular encephalopathy, hypertension, hypercholesterolemia, AD associated depression |
| C1_16 | 76 | male | 12 | 25 | Vitamin D deficiency, positive borrelia-IgG, cerebral microbleeds |
| C1_17 | 69 | female | 11 | 30 | AD associated depression |
| C1_18 | 56 | female | 10 | 19 | None |
| C1_19 | 51 | male | 12 | 20 | AD associated depression |
| C2_01 | 71 | female | 13 | 17 | Hypertension, AD associated depression, hypothyroidism |
| C2_02 | 70 | male | 11 | 21 | Hypertension, diabetes II |
| C2_03 | 74 | male | 18 | 24 | Coronary heart disease, hypertension, hypercholesterolemia, indigestion |
| C2_04 | 78 | male | 20 | 28 | Spinal canal stenosis L4/L5, arterial hypertension, indigestion |
| C2_05 | 66 | male | 14 | 23 | Paroxysmal symptomatic atrial fibrillation |
| C2_06 | 84 | female | 8 | 20 | Arterial hypertension, polyneuropathy, struma nodosa, cardiac decompensation |
| C2_07 | 58 | female | 15 | 19 | None |
| C2_09 | 70 | female | 8 | 23 | Arterial hypertension, hypothyroidism (euthyreote struma) |
| C2_10 | 78 | male | 14 | 18 | Hypertension, benign hallucinations |
| C2_11 | 79 | female | 8 | 22 | Hypertension, hypothyroidism |
| C2_12 | 76 | female | 15 | 20 | Hypertension, hyperuricemia |
| C2_13 | 59 | male | 20 | 29 | None |
| C2_14 | 72 | female | 9 | 22 | Hypertension, hypercholesterolemia |
| C2_15 | 80 | male | 12 | 27 | Diabetes II, hypertension, hypothyroidism |
| C2_16 | 71 | female | 19 | 29 | Intermittent spinal complaints |
| C2_17 | 72 | male | 12 | 21 | Coronary heart disease, peripheral arterial occlusive disease (legs) |

##### Common Inclusion / Exclusion Criteria

Inclusion criteria:

– Clinically stable patients with probable AD (Diagnosis according to ICD-10 (F00) and NIA-AA Criteria by an expert in cognitive neurology)
– At least 3 months of stable anti-dementia therapy or no anti-dementia therapy necessary
– Signed written informed consent
– Age ≥ 18 years

Exclusion criteria:

– Non-compliance with the protocol
– Relevant intracerebral pathology unrelated to the AD (e.g. brain tumor)
– Hemophilia or other blood clotting disorders or thrombosis
– Corticosteroid treatment within the last 6 weeks before first treatment
– Pregnant or breastfeeding women

#### 2.4.2. Brain stimulation procedure

Since neurodegeneration in AD brains is extended and promising animal data for whole brain ultrasound therapy in Alzheimer’s mouse models exist (Eguchi et al. 2018), we compared non-navigated global cortical stimulation (center 2) with navigated AD network stimulation (center 1). Center 1 used regions of interest (ROI) that were ellipsoids defining the stimulated brain area which should precisely be targeted (requiring a highly focal technique, Figure 6). TPS was performed with single ultrasound pressure pulses: duration about 3 μs (see Figure 1), 0.2 mJ/mm2 energy flux density, pulse repetition frequency 5 Hz, pulse number per therapeutic session 6000. A NEUROLITH TPS generator (Storz Medical AG, Tägerwilen, Switzerland) was utilized. At center 1, a system with navigation capabilities via a visualization tool was applied. The system is CE approved for AD treatment and equipped with an infrared camera system (Polaris Vicra System by Northern Digital Inc.). The camera tracks the positions of the handpiece and the head of the patient (via goggles affixed with infrared markers). To standardize treatments for all patients by means of treatment visualization and recording, the system allows defining standardized target volumes of interest for each individual participant’s MRI (ROIs). This individual tracking enables standardized focal brain stimulation over the whole study population with adequate movements of the handpiece over the skull. Additionally, tracking of the handpiece movement is possible with each pulse leaving a colored mark in the visualization. The whole treatment session was recorded for post hoc evaluation of the hand piece movement and regarding the distribution of pulses for each patient. Depth regulation of the stimulation focus was achieved via variable stand offs at the handpiece. The treatment comprised 3 sessions per week for 2-4 weeks.

**Fig. 6.**
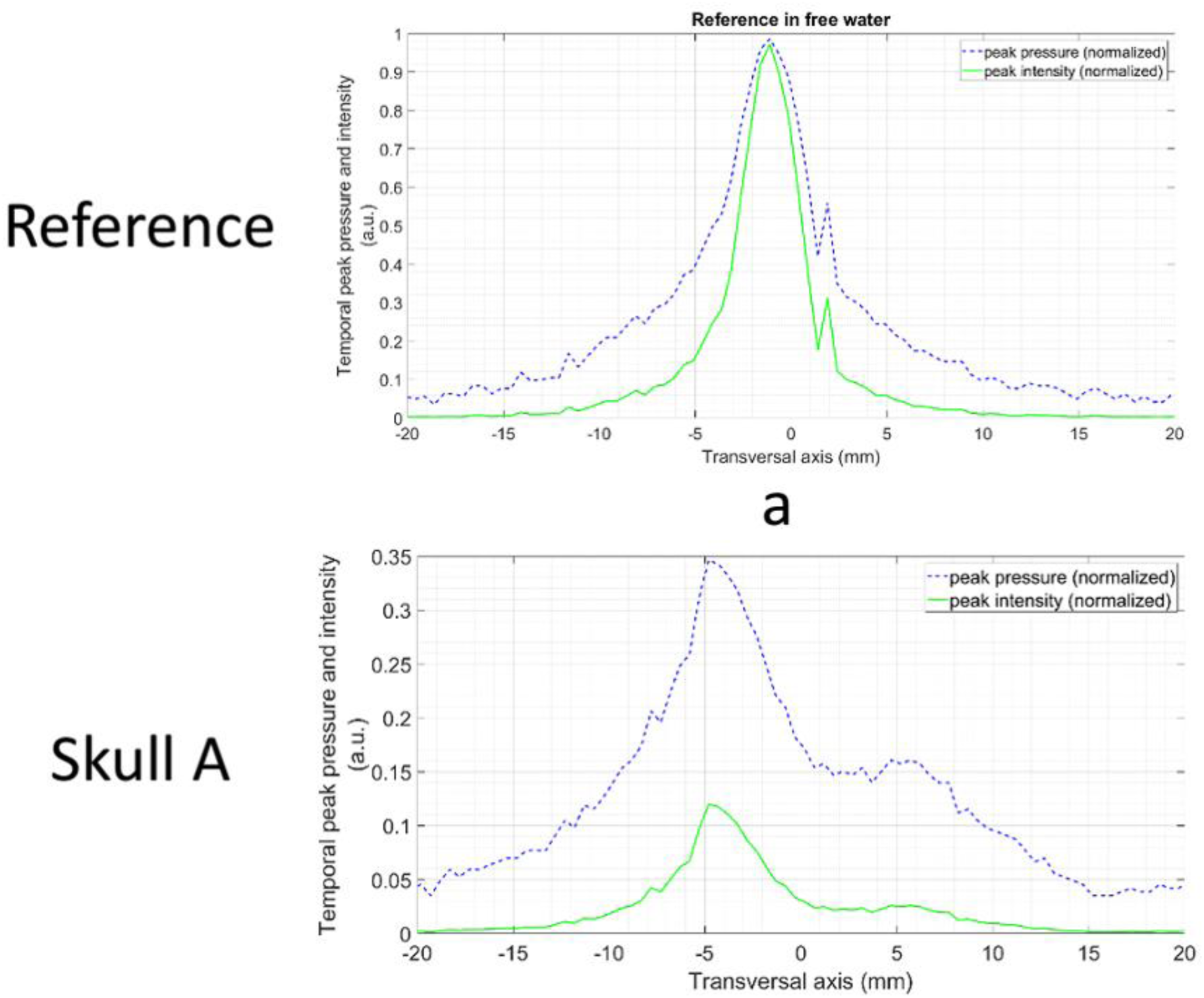
Normalized measured temporal peak pressure and intensity distributions for free water reference and skull A. Y-axis values for skull A are relative to the water reference. The data show the focality of the pulse (= lateral spatial resolution) in the transversal plane measured at beam axis maximum (compare Figure 3).

##### Detailed stimulation procedure at center 1

Individual ROIs were defined by a neurologist to target AD relevant brain areas (AD network). ROIs included the classical AD stimulation target dorsolateral prefrontal cortex and areas of the memory (including default mode) and language networks. According to an anatomical pre-evaluation of brain size variability (in house software for gross estimation of cerebrum size), 2 sets of standardized ROI sizes were established and applied for either small or large patient brains. Every ROI was stimulated twice per session and most patients were stimulated for 4 weeks. Individually set ROIs comprised: bilateral frontal cortex (dorsolateral prefrontal cortex and inferior frontal cortex extending to Broca’s area, ROI volume 136/164 cm³ - 2*800 pulses per hemisphere), bilateral lateral parietal cortex (extending to Wernicke’s area, ROI volume 122/147 cm³ – 2*400 pulses per hemisphere) and extended precuneus cortex (1 bilateral volume with 66/92 cm³ - 2*600 pulses). The goal was to distribute all pulses within the respective ROIs with a focus on the cortical tissue (Figure 1).

##### Stimulation procedure center 2

A non-standardized and non-navigated global brain stimulation approach was performed to compare different modes of TPS stimulation (for comparison see Eguchi et al. 2018). Here, the goal was to homogenously distribute the total energy of 6000 TPS pulses per session over all accessible brain areas over a treatment period of 2 weeks. For this, the TPS handpiece was moved along the anterior-posterior skull axis over the whole scalp as well as in circular motions around the head.

#### 2.4.3. Neuropsychological evaluation

Neuropsychological tests were performed before the stimulation (baseline), after 1-3 days (post-stim), 1 month (1month post-stim), and 3 months after the last stimulation session (3month post-stim).

##### CERAD

The German version of the CERAD Plus (including Trail Making Test and Phonemic word fluency) was used for testing neuropsychological functions. The CERAD is well suited for mild dementia evaluations since it does not show repetition effects with mild AD (Camargo et al. 2015, Matthews et al. 2013). Additionally, the CERAD highly correlates with other global cognitive and functional scales including ADAS-COG (Seo et al. 2010). Evaluations include word fluency (phonemic and categorical), naming (Boston Naming Test), encoding, recognition, and recall of verbal material (Word list), as well as constructional praxis and constructional recall (Figures Copy and Recall). The Trail-Making-Test was accomplishable in only about half of the patients and was thus excluded from final analyses. The CERAD raw scores were used to calculate the Corrected Total Score (CTS, Chandler et al. 2005; N = 35 complete datasets). The Logistic Regression score (LR, Ehrensperger et al. 2010; N = 31) and the principle component analysis scores (PCA, N = 30) were generated using the z-transformed scores (corrected for age, gender, and formal education as performed by the CERAD Online analysis; norm population CERAD: N = 1,100, phonemic word fluency: N = 604). The LR score weights those CERAD subtests which are particularly indicative of AD type dementia (for comparison see the long-term study by Bessi et al. 2018). The PCA on all CERAD subtests defined statistically independent factors that explained individual test performance with an eigenvalue > 1. This approach is similar to the PCA approach by Ehrensperger et al. 2010, but it additionally includes the phonemic word fluency test. For the PCA, the rotation method Varimax with Kaiser normalization was used (SPSS v24).

##### Assessments of depressive symptoms

As depression is a typical comorbidity of AD, we monitored effects on depressive symptoms with the Geriatric Depression Scale (GDS, 30 complete patient datasets) and the Beck Depression Inventory (BDI, 25 complete patient datasets). As GDS and BDI values were not normally distributed according to the Kolmogorow-Smirnow-Test, statistical evaluation was performed with the nonparametric Friedman-Test for multiple paired variables (SPSS v24).

##### Statistical analysis

Statistical analysis of the dependent variables (CERAD CTS, LR score, PCA factors) was done with SPSS v24 applying a mixed ANOVA with TIME as within-subject factor (baseline, post-stim, 1month post-stim, 3months post-stim) and CENTER (1, 2) as between-subject factor. Spearman rank correlation analysis was used to evaluate correlations between the CERAD variables and the depression scores (GDS, BDI).

#### 2.4.4. Functional imaging performed at center 1

At center 1, all patients underwent MR investigations (3 T SIEMENS PRISMA MR with a 64-channel head coil) including anatomical scans used for navigation via the tracking and visualization tool (N = 19 complete datasets). In addition, resting state scans (N = 18) as well as T2* and FLASH images for safety evaluations (bleedings, edema, morphology) were recorded before and after the stimulation interventions.

##### Functional MRI sequence parameters

We recorded a T1-weighted structural image using a MPRAGE sequence (TE/TR = 2.7/1800ms, inversion time = 900ms, flip angle = 9°, resolution 1 mm isotropic). For functional images we used a T_2_*-weighted gradient-echo-planar imaging (EPI) sequence, with 38 slices aligned to AC-PC, covering the whole brain including cerebellum (TE/TR = 30/2500 ms, flip angle = 90°, in-plane acceleration = GRAPPA 2, field of view = 230 x 230 mm, voxel size = 1.8 x 1.8 x 3 mm, 25 % gap). For resting state fMRI 250 volumes (10 min 25 s) were recorded with eyes opened and use of a fixation cross.

##### Resting state fMRI analysis

All preprocessing procedures and analyses were performed using the CONN toolbox v17 (Whitfield-Gabrieli and Nieto-Castanon 2012). This included default preprocessing: realignment, unwarping, slice-time correction, structural segmentation, normalization, outlier detection (ART-based scrubbing), and smoothing (8 mm FWHM kernel). Denoising was done using a band pass filter [0.008 to 0.09 Hz], removal of motion confounds (6 motion parameters and their first derivatives), definition of 5 PCA components extracted from the cerebrospinal fluid and the white matter masks (aCompCor, Behzadi et al. 2007) and scrubbing. For first level analysis, a bivariate correlation of the corrected time series of all voxels was calculated.

##### Network definitions for the resting state analysis

Networks investigated comprise predefined networks of the CONN-Toolbox derived from ICA analyses of the HCP dataset (497 subjects) related to cognitive functions, a memory network (literature based selection of regions) and a network of the stimulated areas:

1. Default mode network (CONN): medial prefrontal cortex, bilateral lateral parietal cortex, precuneus
2. Salience network (CONN): anterior cingulate cortex, bilateral anterior insula, bilateral rostral prefrontal cortex, bilateral supramarginal gyrus
3. Dorsal Attention network (CONN): bilateral frontal eye field, bilateral inferior parietal cortex
4. Fronto-Parietal or Central Executive network (CONN): bilateral lateral prefrontal cortex, bilateral posterior parietal cortex
5. Language network (CONN): bilateral inferior frontal gyrus, bilateral posterior superior temporal gyrus
6. Memory network (Benoit and Schacter 2015): bilateral hippocampus and bilateral anterior and posterior parahippocampal cortex plus the default mode network as defined in (1)
7. Network of brain stimulation sites (Center 1): bilateral inferior and middle frontal gyri, bilateral inferior (supramarginal and angular gyri, parietal operculum) and superior parietal cortex, precuneus

##### Second level functional connectivity analysis

For this, paired t-tests between baseline and post-stim functional connectivity (FC) were calculated (correlation of the time series in the ROIs, 0.05 FDR seed-level correction, two-sided) on group level.

##### Second level connectivity analysis with graph theoretical approach

The graph theoretical measure global efficiency (GE) as an estimate of the capacity of parallel information processing within a network (Achard and Bullmore, 2007) was calculated and compared between the baseline and the post-stim session (adjacency matrix threshold: correlation coefficient = 0.35 to define connected ROIs; analysis threshold 0.05 FDR corr., paired t-tests, two-sided).

##### Correlation of graph theoretical values with neuropsychological scores

Global efficiency (GE) values for networks showing a significant difference between sessions were extracted for all subjects and sessions and were used for correlation analysis with CERAD scores (CTS, LR, PCA factors). Data of both MR sessions entered the correlation analysis which was performed with SPSS 24 using a non-parametric Spearman rank correlation analysis as GE values were not normally distributed.

#### 2.4.5. Patient evaluations

At both centers, patient evaluations were performed at each visit (clinical examinations and patient reports). Center 1 used additional questionnaires to acquire further quantitative information on patients’ treatment experience, including improved functionality and tolerability. These included the German SEG scale (“Skala zur Erfassung der Gedächtnisleistung” = scale for subjective evaluation of memory performance), Inventory of activities of daily living (IADL) questionnaire and a German scale for leisure activity (“Freizeitverhalten”). These questionnaires were applied at all 4 time points of neuropsychological testing (baseline, post-stim, 1month post-stim, 3months post-stim).

In addition, after each of the treatment sessions patients evaluated their pain and pressure experience during treatment (visual analogue scales, 0 = none, 10 = very strong pain/pressure). Patients also reported on side effects with non-standardized answers. For cognitive changes, changes of general activity, mood changes and “change of body state” (control question) patients’ answers were categorized in “improved”, “stable”, and “worsened”.

## 3. Results

### 3.1. Focal TPS energy transmission

#### 3.1.1. TPS data simulations

3D simulation of temporal peak intensities showed that a highly focal energy pulse can be generated through the skull. Calculations for 2 human skulls showed a consistent peak intensity drop (skull attenuation) of about 65% at spatial peak (Figure 3a).

#### 3.1.2. Human skull and brain sample measurements

Measurements of temporal peak intensities with 2 human skulls and 10 human brain samples confirmed the transmission of a focal energy pulse without occurrence of secondary maxima. However, compared to free degassed water, the human skulls produced a temporal-peak intensity drop of 80-90% (Figure 3b). For human brain tissue we found a considerable variability of results depending on the state of the post mortem tissue and the suspected amount of decay gases (brains were 0-7 days post mortem). With consideration of tissue state, overall results corresponded to published values for sound wave attenuation in human brain tissue, which is in the range of 0.58 dB/cm/MHz (Szabo et al. 2014).

#### 3.1.3. Rat skull measurements

Rat skull measurements (Figures 4 and 5) again confirmed a focal energy pulse. Rat skulls produced a mean pressure drop of about 29%. Depending on pulse energy, the detailed losses were: 20.3% for 0.1 mJ/mm^2^, 28.8% for 0.35 mJ/mm^2^ and 37.3% for 0.55 mJ/mm^2^.

Overall, spatial-peak-temporal-average intensities (Ispta) are in the same range as published data of LIFUS technologies. However, spatial-peak-pulse-average intensities (Isppa) are higher with TPS compared to LIFUS values.

### 3.2. TPS safety

#### 3.2.1. In vivo evaluations of 80 rats

80 rats were treated with a constant TPS focus within the brain for a TPS safety investigation (energy levels 0-0.3 mJ/mm2, 100-400 focal pulses; Ameln-Mayrhofer, in preparation). Brain preparations did not show any intracranial, subarachnoid or subdural bleeding in any rat of any group. Histological investigations of two brains per group did not show any abnormalities in any TPS group. Despite considerably lower pressure wave attenuation by rat skulls (29% in rat instead of 85% in human skulls), TPS application in living rats was safe – even at higher energy levels than those used for the clinical study.

#### 3.2.2. Patient evaluations

At both centers patient evaluations at follow ups up to 3 months (clinical examinations and patient reports) did not show any relevant side effects (Table 5). A detailed quantification performed by center 1 resulted in 4% headache (partly previous headache history present), 3% mood deterioration, and 93% none. Visual analogue scale evaluation (0-10) of within-treatment pain or pressure experience resulted in 92% 0, 7% 1-5, 1% 6-8 (pain) and 83% 0, 15% 1-5, 2% 6-8 (pressure).

#### 3.2.3. MRI evaluations

Evaluation of anatomical MRIs as well as T2* and FLASH images before and after stimulation did not reveal any hemorrhages, edema or any other type of new intracranial pathology.

### 3.3. Neuropsychological improvements in Alzheimer’s Disease

#### 3.3.1 CERAD CTS

CERAD CTS score analysis (N = 35) revealed a significant within-subjects effect of TIME: *P* < .0001. The between-subjects effect of CENTER was not significant (*P* = .313). This indicates that CTS values differ between the 4 time points, but not overall between centers. Post-hoc pairwise comparisons of CTS values (Bonferroni corrected) unveil significant differences for the following: baseline < post-stim (*P _Bonf_* < .0001), baseline < 1month post-stim (*P _Bonf_* < .0001), baseline < 3months post-stim (*P _Bonf_* < .0001; see Table 3 and Figure 7a). Furthermore, a significant interaction TIME*CENTER (*P* = .003) was found indicating that CTS differences between time points vary between the centers. A follow-up repeated measurements ANOVA for both centers separately revealed a significant main effect of TIME for both centers individually. All 3 pairwise comparisons remained significant for center 2, whereas for center 1 only the baseline < post-stim contrast reached significance.

**Fig. 7.**
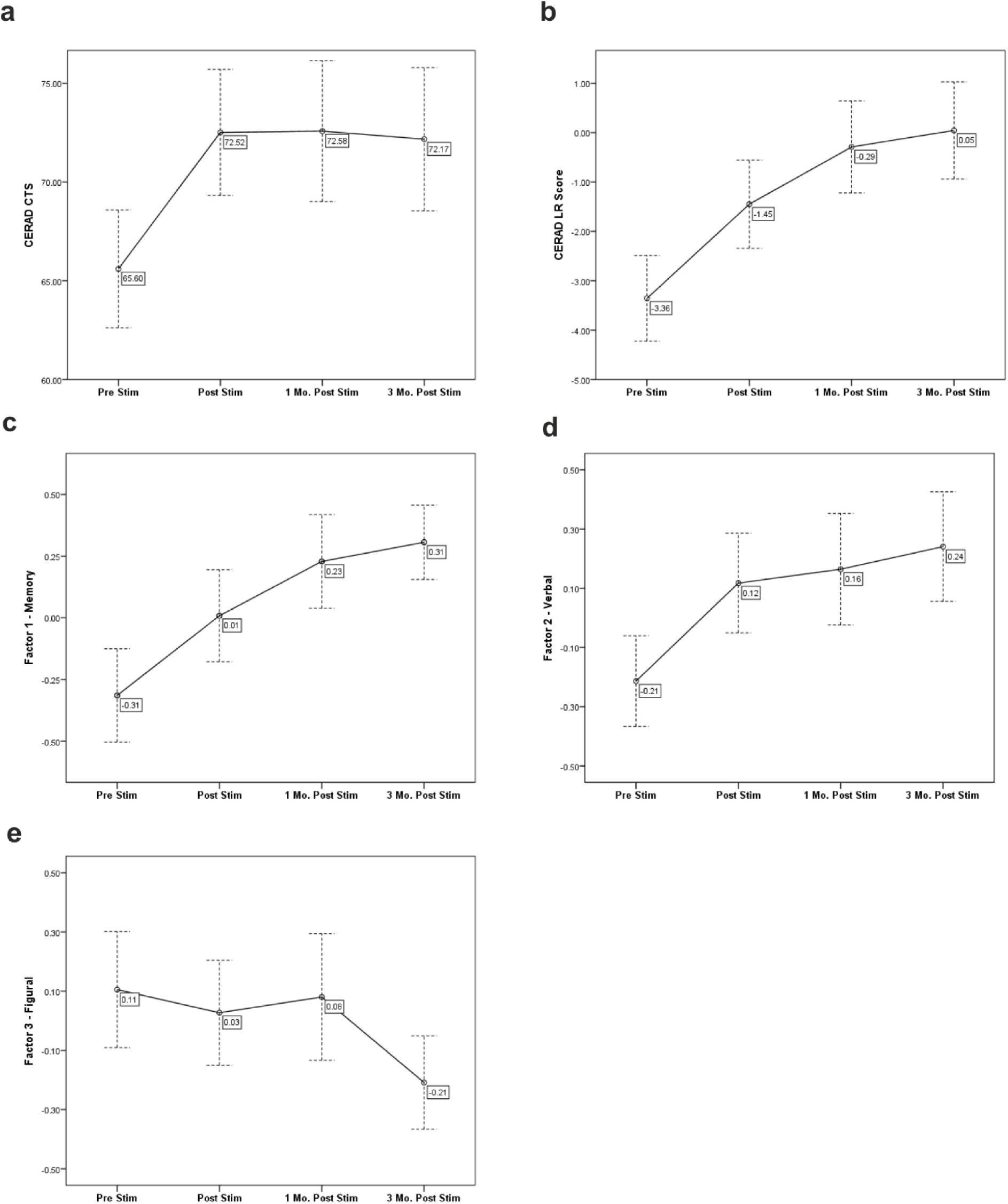
CERAD score changes (mean +/-1 standard error) over time. **a-b,** The average CERAD summary scores (CTS and LR scores) show a significant increase over time. **c-e,** The first CERAD factor loadings representing Memory (Factor 1, **c**) und Verbal functions (Factor 2, **d**) increase over time while for Factor 3 (Figural functions, **e**) a decline was observable

**Table 3.** Results of the mixed ANOVA for the combined dataset with post-hoc comparisons between time points

| | N | Factor | F | p | $\eta_p^{2*}$ | Sign. Contrasts (P)** |
| --- | --- | --- | --- | --- | --- | --- |
| **CTS** | 35 | Time | $F_{(3,99)} = 18.304$ | .000 | .357 | 1 – 2 (.000)<br>1 – 3 (.000)<br>1 – 4 (.000) |
| | | Center | $F_{(1,33)} = 1.051$ | .313 | .031 | |
| | | Time*Center | $F_{(3,99)} = 4.892$ | .003 | .129 | |
| **LR** | 31 | Time | $F_{(2,69)} = 20.129$<br>(GG) | .000 | .410 | 1 – 2 (.000)<br>1 – 3 (.000)<br>1 – 4 (.000)<br>2 – 3 (.012) |
| | | Center | $F_{(1,29)} = .047$ | .830 | .002 | |
| | | Time*Center | $F_{(2,69)} = 3.220$<br>(GG) | .038 | .100 | |
| **Factor 1**<br>**(Memory)** | 30 | Time | $F_{(3,84)} = 7.050$ | .000 | .201 | 1 – 3 (.002)<br>1 – 4 (.002) |
| | | Center | $F_{(1,28)} = .272$ | .606 | .010 | |
| | | Time*Center | $F_{(3,84)} = .828$ | .482 | .029 | |
| **Factor 2**<br>**(Verbal)** | 30 | Time | $F_{(3,84)} = 12.433$ | .000 | .307 | 1 – 2 (.003)<br>1 – 3 (.001)<br>1 – 4 (.000) |
| | | Center | $F_{(1,28)} = 2.339$ | .137 | .077 | |
| | | Time*Center | $F_{(3,84)} = 5.351$ | .002 | .160 | |
| **Factor 3**<br>**(Figural)** | 30 | Time | $F_{(3,84)} = 3.739$ | .014 | .118 | 1 – 4 (.007) |
| | | Center | $F_{(1,28)} = 2.033$ | .165 | .068 | |
| | | Time*Center | $F_{(3,84)} = 3.708$ | .015 | .117 | |

\* Partial eta squared as an estimate for the effect size.
\*\*Post-hoc comparisons between time points (1 = baseline, 2 = post-stim, 3 = 1month post-stim, 4 = 3months post-stim) were corrected for multiple comparisons using Bonferroni correction.
CTS = CERAD Corrected Total Score, LR = Logistic regression score, GG = Greenhouse-Geisser corrected.

#### 3.3.2 CERAD LR score

For the CERAD LR score (N = 31) a significant within-subjects effect of TIME (*P* < .0001, Greenhouse-Geisser corrected, since the assumption of sphericity was violated) was found. In contrast, the between-subjects effect of CENTER was not significant (*P* = .830) indicating that LR values differ among the 4 time points but not between the centers. Post-hoc pairwise comparisons revealed significant differences for baseline < post-stim (*P _Bonf_* < .0001), baseline < 1month post-stim (*P _Bonf_* < .0001), baseline < 3months post-stim (*P _Bonf_* < .0001), post-stim < 1month post-stim (*P _Bonf_* = .012; Table 3, Fig. 7b). As for the CTS score, a significant interaction TIME*CENTER (*P* = .038) was found. Further, the main effect of TIME was significant for both centers in a repeated measurements ANOVA for both centers separately. Again, for center 2 all 3 baseline comparisons remained significant, but for center 1 only the baseline < 1month post-stim comparison reached significance.

#### 3.3.3. CERAD PCA

Three factors achieved eigenvalues greater than 1 which means that they explained more variance than every single subtest taken alone (Table 4). Factor 1 (eigenvalue = 5.09, explained variance = 46.25%) displayed the highest loadings on the delayed recall and recognition of the Word List and on Savings of the Word List and the Figures and was thus named Factor “Memory”. Factor 2 (eigenvalue = 1.53, explained variance = 13.95%) was interpreted as “Verbal functions” as its highest loadings were found for the Verbal Fluency tasks and the Word List Total. The loadings of Factor 3 (eigenvalue = 1.19, explained variance = 10.77%) were highest for the figural tasks and this factor was termed “Figural” functions.

**Table 4:** PCA factor loadings of the CERAD subtests.

| CERAD variables | Factors* |  |  |
| --- | --- | --- | --- |
|  | 1<br>Memory | 2<br>Verbal | 3<br>Figural |
| Animal Fluency | .118 | **.872** | .306 |
| Boston Naming Test | .312 | **.478** | .146 |
| Word List – Total | .524 | **.702** | .199 |
| Word List – Delayed Recall | **.834** | .395 | .159 |
| Word List – Intrusions | **.579** | .204 | .004 |
| Word List – Savings | **.819** | -.070 | .073 |
| Word List – Recognition | **.649** | .482 | .004 |
| Figures – Copy | .005 | .121 | **.922** |
| Figures – Delayed Recall | .583 | .118 | **.720** |
| Figures – Savings | **.734** | .172 | .268 |
| Phonemic Fluency | .038 | **.895** | -.113 |

\* Factors with an eigenvalue > 1 derived from a principle component analysis (PCA) with the rotation method Varimax with Kaiser normalization. Factors were interpreted as “Memory” (Factor 1), “Verbal functions” (Factor 2), and “Figural functions (Factor 3) according to highest loadings of the CERAD variables on the three factors (bold).

##### Factor 1 (MEMORY)

A mixed ANOVA with the PCA factor loadings (N = 30) showed a significant within-subjects effect of TIME: *P* < .0001 (Table 3, Figure 7c). The between-subjects effect of CENTER (*P* = .606) as well as the interaction TIME*CENTER (*P* = .482) were not significant. Post-hoc pairwise comparisons showed significant differences for baseline < 1month post-stim (*P _Bonf_* = .002) and baseline < 3months post-stim (*P _Bonf_* = .002).

##### Factor 2 (VERBAL)

Again, the mixed ANOVA showed a significant within-subjects effect of TIME: *P* < .0001 (Table 3, Figure 7d). The between-subjects effect of CENTER was not significant (*p* = .137). Post-hoc pairwise comparisons demonstrated significant differences for baseline < post-stim (*P _Bonf_* = .003), baseline < 1month post-stim (*P _Bonf_* = .001), baseline < 3months post-stim (*P _Bonf_* < .0001). Further, a significant interaction TIME*CENTER (*P* = .002) was found indicating that factor differences between time points vary between centers. The main effect of TIME was significant for center 2 only (repeated measurements ANOVA for the centers separately) and all 3 pairwise comparisons remained significant.

##### Factor 3 (FIGURAL)

Factor 3 revealed a significant within-subjects effect of time in the sense of a DECLINE (*P* = .014; Table 3, Figure 7e). The between-subjects effect of CENTER was not significant (*P* = .165). Post-hoc pairwise comparisons showed a significant decline for baseline > 3month post-stim (*P _Bonf_* = .007). In addition, a significant interaction TIME*CENTER (*P* = .015) was found demonstrating that factor differences between time points differ between centers. The main effect of TIME was significant for center 1 only and the decline for baseline > 3month post-stim remained significant (repeated measurements ANOVA for the centers separately). A further qualitative evaluation indicates that this effect is primarily due to a considerable decline of constructional praxis (Figures – Copy) of the patients of center 1.

#### 3.3.4. Depression scores cannot explain neuropsychological improvement

For the GDS, the effect of TIME was significant (*P* = .005). Pairwise comparisons (Wilcoxon-Test) showed GDS improvement for baseline > 3months post-stim (*P_Bonf_* = .012). For BDI, effect of TIME was also significant (*P* < .0001). Pairwise comparisons (Wilcoxon-Test) displayed BDI improvement for baseline > post-stim (*P_Bonf_* = .012), baseline > 1month post-stim (*P_Bonf_* = .006) and baseline > 3months post-stim (*P_Bonf_* = .012). Importantly, there was no significant correlation between BDI / GDS scores and global CERAD scores (CTS, LR) or the PCA factors after accounting for multiple comparisons (Bonferroni correction). This indicates that CERAD improvements were not driven by changes of depressive symptoms.

#### 3.3.5. Improvement of subjective patient performance

Results from post treatment standard scales showed significant improvements in the subjective evaluation of memory performance (SEG) over time (within-subjects effect of time: *P* = .027, pairwise comparisons not significant). The other standard scales did not show significant changes. In the post-treatment questionnaires, up to 20% of the patients reported subjective improvements and only 2-3% aggravations (details in Table 5).

**Table 5.** Results from post treatment patient questionnaires

|  | Worsened *in* % | Stable *in* % | Improved *in* % |
| --- | --- | --- | --- |
| Cognition | 2 | 77 | 20 |
| General activity | 3 | 82 | 15 |
| Mood | 3 | 71 | 26 |
| Body state* | 2 | 93 | 5 |

\* Questions regarding the body state represent a control item as general somatic changes were not assumed to be associated with the TPS intervention. Post treatment questionnaires were acquired at Center 1 only (N = 19).

### 3.4. Improved memory network connectivity

In the functional connectivity (FC) analysis only the memory network showed significantly increased connectivity when comparing baseline vs. post-stim. Specifically, an increased interhemispheric connectivity was found: the left parahippocampal cortex (anterior part) showed a significant connectivity increase to the right parahippocampal cortex (posterior part). For all other networks, baseline vs. post-stim FC contrasts were not significant (FDR 0.05 corrected, two-sided).

In the graph theoretical analysis again only the memory network showed significant differences between sessions (GE values, post-stim > baseline; Fig. 8). This was found on network level and on ROI level. The following memory network ROIs showed significant GE value increases: hippocampus (bilateral), left parahippocampal cortex (anterior and posterior), left lateral parietal cortex, right parahippocampal cortex (anterior), precuneus. No other network investigated displayed a significant difference between the sessions. Correlating all baseline and post-stim GE values with the CERAD scores, the CTS, the LR and the Memory factor scores generated significant results, which indicates that the memory network GE values are indeed related to cognitive performance (Table 6).

**Fig. 8.**
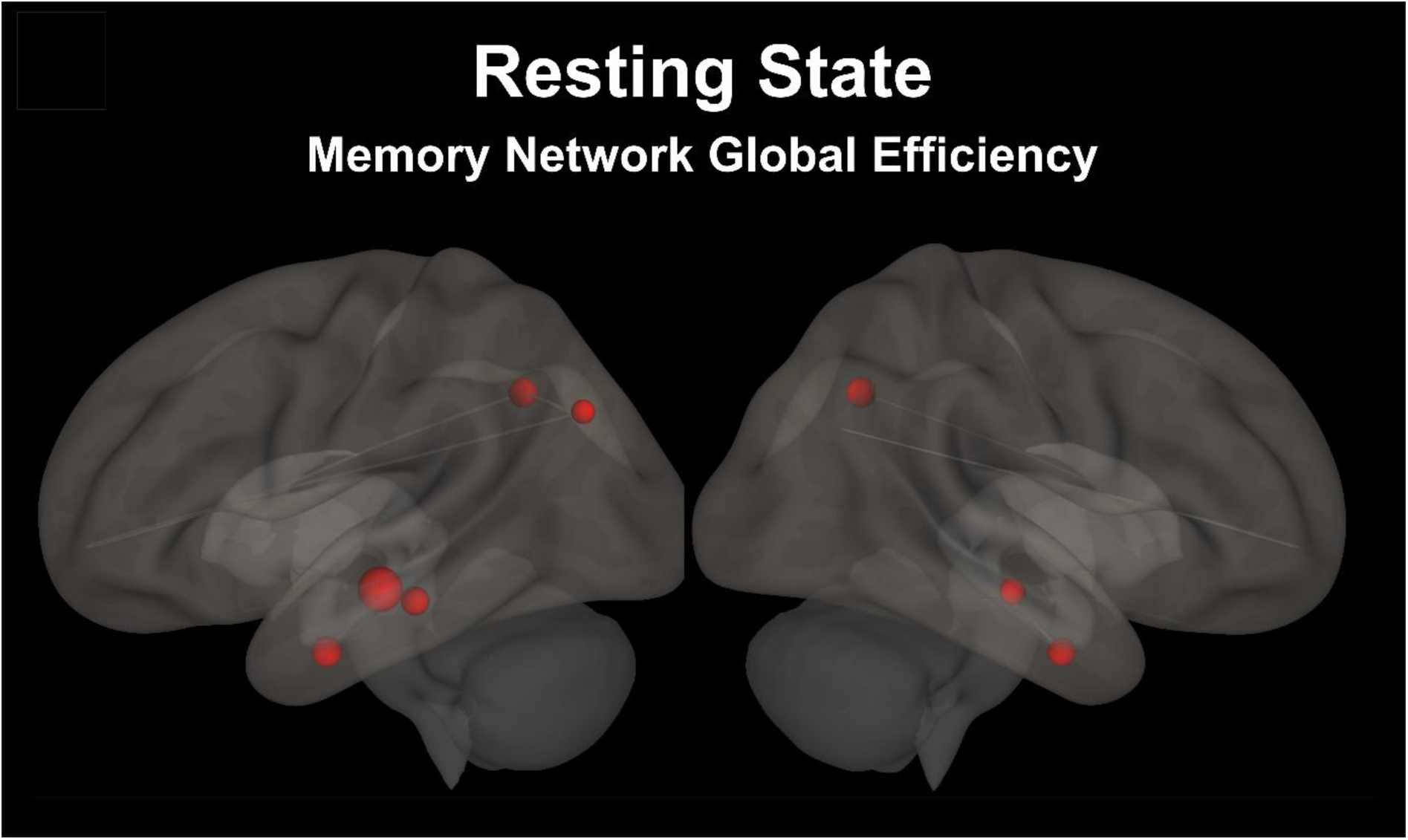
Increased functional connectivity in the memory network after the stimulation. Graph theoretical analysis of the global efficiency (GE) of the memory network during the resting state scan (18 patients) demonstrated significantly increased global efficiency after the stimulation in bilateral hippocampal and parietal areas of the memory network as indicated by the red spheres (sphere size weighted according to GE values).

**Table 6.**
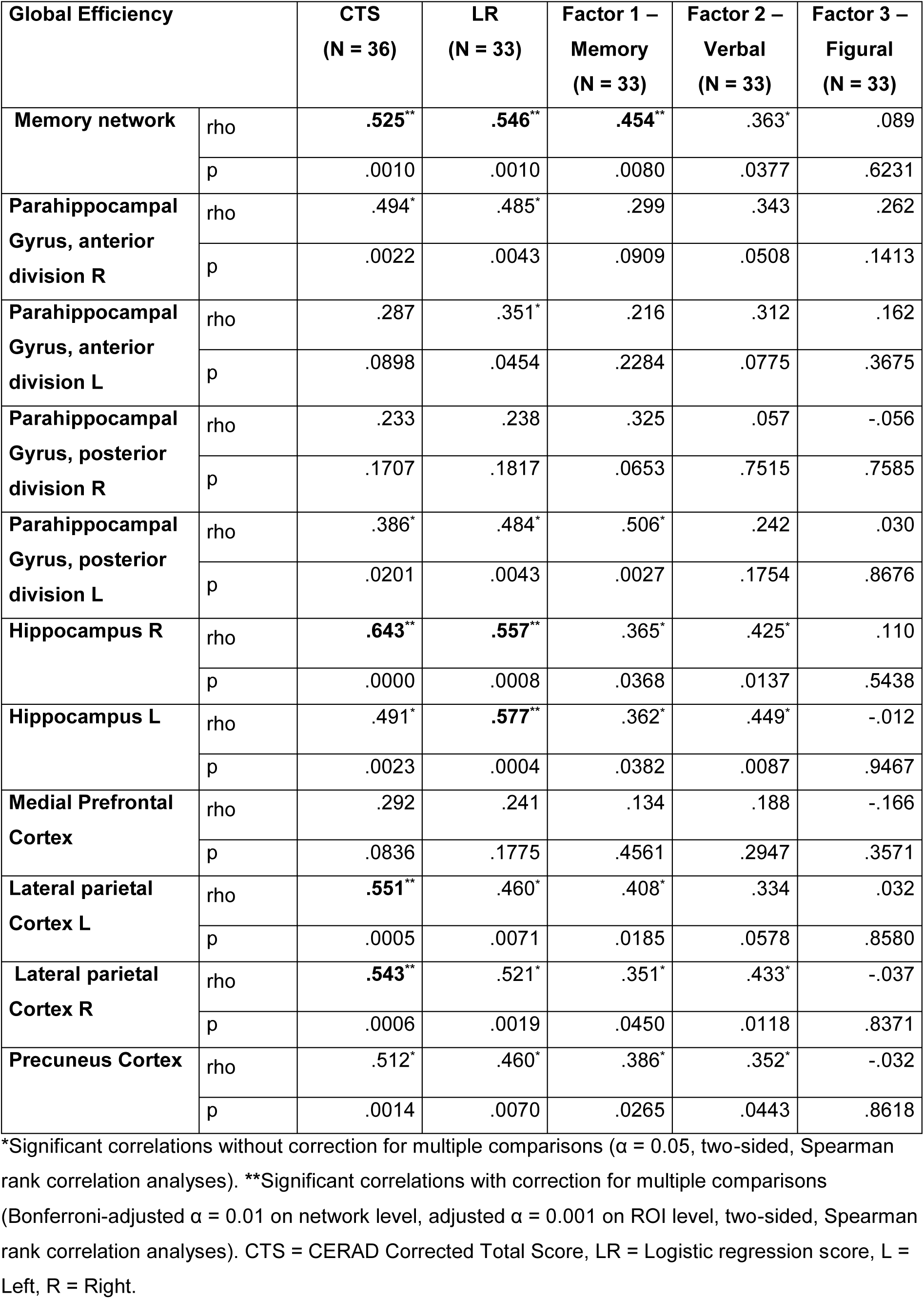
Correlation between the resting state global efficiency values in the memory network and its nodes, with the neuropsychological test scores.

| Global Efficiency |  | CTS<br>(N = 36) | LR<br>(N = 33) | Factor 1 –<br>Memory<br>(N = 33) | Factor 2 –<br>Verbal<br>(N = 33) | Factor 3 –<br>Figural<br>(N = 33) |
| --- | --- | --- | --- | --- | --- | --- |
| Memory network | rho | .525** | .546** | .454** | .363* | .089 |
|  | p | .0010 | .0010 | .0080 | .0377 | .6231 |
| Parahippocampal<br>Gyrus, anterior<br>division R | rho | .494* | .485* | .299 | .343 | .262 |
|  | p | .0022 | .0043 | .0909 | .0508 | .1413 |
| Parahippocampal<br>Gyrus, anterior<br>division L | rho | .287 | .351* | .216 | .312 | .162 |
|  | p | .0898 | .0454 | .2284 | .0775 | .3675 |
| Parahippocampal<br>Gyrus, posterior<br>division R | rho | .233 | .238 | .325 | .057 | -.056 |
|  | p | .1707 | .1817 | .0653 | .7515 | .7585 |
| Parahippocampal<br>Gyrus, posterior<br>division L | rho | .386* | .484* | .506* | .242 | .030 |
|  | p | .0201 | .0043 | .0027 | .1754 | .8676 |
| Hippocampus R | rho | .643** | .557** | .365* | .425* | .110 |
|  | p | .0000 | .0008 | .0368 | .0137 | .5438 |
| Hippocampus L | rho | .491* | .577** | .362* | .449* | -.012 |
|  | p | .0023 | .0004 | .0382 | .0087 | .9467 |
| Medial Prefrontal<br>Cortex | rho | .292 | .241 | .134 | .188 | -.166 |
|  | p | .0836 | .1775 | .4561 | .2947 | .3571 |
| Lateral parietal<br>Cortex L | rho | .551** | .460* | .408* | .334 | .032 |
|  | p | .0005 | .0071 | .0185 | .0578 | .8580 |
| Lateral parietal<br>Cortex R | rho | .543** | .521* | .351* | .433* | -.037 |
|  | p | .0006 | .0019 | .0450 | .0118 | .8371 |
| Precuneus Cortex | rho | .512* | .460* | .386* | .352* | -.032 |
|  | p | .0014 | .0070 | .0265 | .0443 | .8618 |

\*Significant correlations without correction for multiple comparisons ( $\alpha = 0.05$ , two-sided, Spearman rank correlation analyses). \*\*Significant correlations with correction for multiple comparisons (Bonferroni-adjusted $\alpha = 0.01$ on network level, adjusted $\alpha = 0.001$ on ROI level, two-sided, Spearman rank correlation analyses). CTS = CERAD Corrected Total Score, LR = Logistic regression score, L = Left, R = Right.

## 4. Discussion

The application of ultrasound for brain therapy has become a hot topic as it bears the potential for providing a new class of invasive (ablation, e.g. Lee et al. 2019, Martinez-Fernandéz et al. 2018, Elias et al. 2016), semi-invasive (blood brain barrier opening, e.g. Lipsman et al. 2018) or non-invasive (neuromodulation, e.g. Lohse-Busch et al. 2014, Monti et al. 2016) brain therapies. In this manuscript, we describe a new non-invasive technique, Transcranial Pulse Stimulation (TPS), and we provide first clinical data indicating that the method is safe and clinically feasible. Treatment tolerability in patients was high and no relevant side effects were noted. The in vivo animal study tested 6-7 fold higher energy levels as compared to our clinical pilot study and did not cause any tissue damage.

In addition to safety and feasibility, preliminary efficacy data in AD patients indicate that the therapy may improve neuropsychological test scores with consequences for the subjective functioning of the patients (subjective memory improvement). Importantly, improvements reported in this study indicate 3 month long-term effects of ultrasound neuromodulation. Note, that effects were achieved in an AD population already receiving optimized standard treatment. Our principle component analysis showed that besides improvements in the language domain, there was a particular improvement of memory performance. Resting-state fMRI analyses revealed an enhanced functional connectivity and global efficiency in the memory network that might drive the behavioral improvements. Both is in line with the well-known cortical stimulation effects on deep network nodes (compare Fox et al. 2014, Wang et al. 2014, Lee et al. 2016).

Regarding the site specific outcome, differences between centers were small despite having used different approaches. This is probably related to the fact, that the navigated and the non-navigated procedure both stimulated large areas of the cortex in a population with widespread pathology. For the figural network, however, an interesting effect was found. Figural tests showed a significant decline after 3 months driven by the constructional praxis results of center 1 only. Center 1 performed a navigated stimulation of the AD network instead of a global brain stimulation and did not comprehensively cover all areas relevant for constructional praxis (e.g. occipito-parietal cortex was not treated). Therefore, a stimulation-site specific effect seems reasonable: at center 1 fMRI and neuropsychological data showed specific upregulation of mnemonic functions only which are supported by the stimulated cortical areas. In contrast, patients figural abilities related to the non-stimulated occipto-parietal areas declined in the tests, which may be compatible with the natural course of the disease.

Concerning the preclinical experiments, results demonstrate that TPS can generate well focused brain stimulation pulses within the brain. The simulations, in vitro and in vivo animal measurements also indicate broad safety margins. Even applying 100 pulses of 0.3 mJ/mm^2^ at the same brain location through rat skulls - in which we measured only about one third of the human skulls attenuation - did not induce any hemorrhage or injury. Note that the patient study applied lower energies at continuously varying locations.

Regarding the precise mechanisms, specifically how ultrasound may affect neurons and generate neuroplastic effects, current knowledge is limited. Several principles, related to different techniques, have been proposed (Tyler et al. 2018, Miller and O’Callaghan 2017, Ventre and Koppes 2016, Bystritsky and Korb 2015). They include pore formations in cell membranes (Babakhanian et al. 2018), increases in extracellular serotonin and dopamine levels (Min et al. 2011), reduction of GABA levels (Yang et al. 2012), increase of brain-derived neurotrophic factor (BDNF), glial cell line-derived neurotrophic factor (GDNF) and vascular endothelial growth factor (VEGF) (Lin et al. 2015). Using ultrashort ultrasound pulses and a neuronal stem cell culture, stem cell proliferation and differentiation to neurons could be enhanced (Zhang et al. 2017). In an AD mouse model, microglia activation with plaque reduction, clearing of Aβ into microglial lysosomes and improvements of spatial and recognition memory has been shown (Leinenga et al. 2015). Another study suggested an important role of nitric oxide synthase when improving cognitive dysfunctions in mouse models of dementia by whole brain stimulation (Eguchi et al. 2018).

Concerning study limitations, this safety and feasibility investigation very likely includes placebo and/or repetition effects and further strict sham-controlled investigations are required to confirm the stimulation effects. However, it seems unlikely that all results are explainable by non-specific effects. First, the long-term course of our neuropsychological improvements clearly differs from the expectable placebo responses as described by Ito et al. (2013). The dissociation between improved memory and language but worsened figural functions is also hard to explain by placebo effects. Second, the fMRI data indicate a predominant improvement of the memory network being compatible with the focused AD network stimulations performed in the fMRI subgroup. For a better assessment of clinical efficacy, follow up studies should compare patient subgroups regarding disease stage and comorbidities. As our primary interest was to judge safety and feasibility on a broad range of patients in a realistic outpatient setting, we did not recruit a homogeneous AD group for this pilot study. In a recent study by Guo et al. (2018), acoustic effects were suggested as possible confounding factors when looking at immediate ultrasound effects on the brain. In our data, there were no indications that simple acoustic effects could be the source of the AD long-term improvements. Importantly, we did not find auditory network changes in the fMRI data, which could have been induced by the ultrasound application. Regarding the optimal mode of TPS application, our experimental design is just an initial step. Although we made sure that energy transmission and safety issues are clarified before starting a patient intervention, future studies should investigate an optimal mode further. The best settings for pulse frequency, pulse intensity, number of sessions and number, location and extension of areas to be stimulated (including subcortical nodes) have yet to be determined. Further, a combination of brain stimulation with cognitive tasks might improve clinical outcome (Rabey and Dobronevsky 2016, Manenti et al. 2018). Also, definitive judgement of safety issues requires data from larger populations. Note however, that our animal data indicate large safety margins and more than 1500 experimental human pilot applications have already been performed over the last years without evidence for major side effects.

Neurodegenerative disorders like Alzheimer’s or Parkinson’s disease are one of the most important medical problems within our ageing society. Available treatments are limited, and such patients are therefore major candidates for clinical benefits of new add on therapies like TPS (Miller et al. 2017). They allow precise targeting of diseased network areas which involves two aspects: (1) stimulation of larger cortical ROIs with well definable stimulation borders and (2) precise stimulation of smaller foci or even deep network nodes. Our preclinical data and the quality of targeting ROI borders demonstrate the focal stimulation potential TPS bears. This is a particular advantage over electrophysiological brain stimulation techniques, where focality and deepness are generally difficult issues, especially in pathological brains (Minjoli et al. 2017).

We conclude that TPS represents a promising novel brain stimulation method with a mobile system adequate for human research and brain stimulation therapy. Results encourage broad neuroscientific application and translation of the new method to clinical therapy and randomized sham-controlled studies.

## Acknowledgements

We are grateful to reviewer comments received on a first description of these data submitted in September 2018. This work was supported by a research-cluster grant from the Medical University of Vienna and University of Vienna (SO10300020) and a research grant from STORZ Medical to R.B. and the Medical University of Vienna (UE76101004). Methodology of the fMRI part was partly developed via support of the Austrian Science Fund FWF (P 23611, KLIF453, KLIF455). Dr. Hallett is supported by the NINDS Intramural program. Mobilitas e.V. (non-profit organization registered in Gemany) supported patient organization in Germany. We acknowledge support in acquisition of the preclinical data by Arvid Kühl, Jürgen Mayer (STORZ Medical) and Wolfgang Weninger, Lukas Reissig (Medical University of Vienna). We acknowledge support in acquisition and analysis of the clinical data by Peter Dal-Bianco, Elisabeth Stögmann, Nina Mahr, Florian Fischmeister, Jakob Rath, Thomas Foki and Eduard Auff.

## Author contributions

General ideas for the experiments were contributed by E.Mar., R.B. (preclinical) and R.B., H.LB., E.Mar (clinical). Design of experiments was done by E.Mar., R.B., C.F., C.G. (preclinical) and R.B., H.LB., E.Mar., C.F., E.M., H.B., J.L. (clinical). Data collection was done by E.Mar., C.G., C.F., R.B. (preclinical) and H.LB., T.PN, H.B., E.M., M.Sch., A.A., T.A., R.R., A.W., R.B., U.R. (clinical). Analysis and interpretation of data was performed by E.Mar., C.G., R.B., C.F. (preclinical) and R.B., E.M., M.Sch., C.F., H.B., H.LB, M.H. (clinical). The manuscript was written by R.B., E.M., C.G., M. Sch., M.H. All authors contributed to editing and revising the manuscript.

